# Intercellular alignment of apical-basal polarity coordinates tissue homeostasis and growth

**DOI:** 10.1101/2021.10.11.463906

**Authors:** Jinghua Gui, Yunxian Huang, Satu-Marja Myllymäki, Marja Mikkola, Osamu Shimmi

## Abstract

Maintaining apicobasal polarity (ABP) is crucial for epithelial integrity and homeostasis during tissue development. Although recent studies have greatly advanced our understanding of intracellular mechanisms underlying ABP establishment, it remains largely unknown how the ABP is regulated at the tissue level. Here, we address intercellular mechanisms coordinating ABP using the *Drosophila* wing imaginal disc. By studying Scribble, a key ABP determinant, we show that ABP is regulated through intercellular alignment, which takes place either progressively or regressively in a context-dependent manner. Cells expressing wild type *scribble* progressively restore ABP in *scribble* hypomorphic mutant cells. In contrast, cells with conditional *scribble* knockdown instigate the regressive loss of polarity in abutting wild type cells. Our data reveal that genetic and physical interactions between Scribble, Septate junction complex and α-Catenin appear to be key for sustaining intercellular network of ABP. Taken together, our findings indicate that the intercellular relay of the status of ABP contributes to the robustness of polarity across the tissue.

## Introduction

In epithelia of developing multicellular organisms, establishing apicobasal polarity (ABP) plays a key role in maintaining tissue function and homeostasis (Bilder, 2004; Hariharan, 2015). Epithelial cells establish distinct sub-compartments along the apicobasal axis and vectorize specific cellular functions (Thompson, 2013). This is achieved through the interaction or antagonism among multiple groups of intracellular polarity determinants and intercellular junctions (Campanale et al., 2017; Thompson, 2013).

Studies employing the *Drosophila melanogaster* wing imaginal disc, a monolayer epithelium comprising columnar cells, have greatly contributed to the understanding of ABP establishment and maintenance. In *Drosophila* epithelial cells, along the apical-to-basal axis lies multiple conserved complexes of polarity determinants: apical complex containing the Crumbs and partitioning defective (PAR) complexes, and basolateral Scribble (Scrib) complex (Bonello and Peifer, 2019; Lang and Munro, 2017; Thompson, 2013). The determinants within the same complex are interdependent as loss of either will dissociate the whole complex. On the other hand, components of distinct complexes are mutually exclusive but interactive, and thus a compromised complex failing to guard the corresponding sub-compartment will lead to either expansion or dismiss of the other complex and eventually complete loss of ABP (Campanale et al., 2017).

The Scrib module, comprising of Scrib, Discs large (Dlg) and Lethal giant larvae (Lgl), localizes at lateral membranes and is enriched at septate junctions (SJs, functional counterpart of mammalian tight junctions (TJs)) (Bilder, 2004; Bonello and Peifer, 2019; Schulte et al., 2006). Loss of the Scrib module often leads to abolishment of ABP, disorganization of tissue architecture, loss of growth control and thus neoplasia (Bilder, 2004; Stephens et al., 2018; Zeitler et al., 2004). This indicates that epithelial cell polarity and growth control are often tightly coupled, reflected in the fact that many molecules identified as ABP determinants serve as tumor suppressors (Bilder, 2004; Hariharan, 2015).

The deregulation of Scrib leads to compromised Hippo signaling pathway (Bonello and Peifer, 2019; Cordenonsi et al., 2011; Yang et al., 2015). The Hippo pathway, a conserved regulator of organ size in animals, was identified first in *Drosophila*, where its loss results in a hippopotamus-shaped phenotype (Hansen et al., 2015; Huang et al., 2005). Hippo signaling is regulated by a variety of upstream signaling pathways including cell polarity regulation, levels of F-actin, tension within the actin cytoskeleton, and cell attachments (Dupont et al., 2011; Fernandez et al., 2011; Gaspar and Tapon, 2014; Hansen et al., 2015; Sansores-Garcia et al., 2011). In *Drosophila*, Yorkie (Yki), the *Drosophila* ortholog of YAP and a key effector of the Hippo pathway, is phosphorylated by Wartz (Wts), a homologue of Large tumor suppressor kinase 1/2 (Lats1/2) in vertebrates, to be retained in the cytoplasm or degraded, otherwise moves into the nucleus to regulate Yki-dependent transcription and growth (Davis and Tapon, 2019; Hansen et al., 2015; Justice et al., 1995).

In epithelial morphogenesis, cell-cell communication through cellular junctions plays a crucial role in tissue development. In both *Drosophila* and mammalian epithelial cells, E-Cadherin-mediated cell-cell adhesion at adherens junctions (AJs) appear to play a central role in mediating mechanical circuit that integrates adhesion, contractile forces and biochemical signaling (Rubsam et al., 2017; Sun and Irvine, 2016). The core components of the AJs are E-Cadherin and their binding partners α-, ß-, and p120-Catenins. E-Cadherin engages in adhesive complexes with neighboring cells, which are connected to the actomyosin cytoskeleton primarily via α-Catenin (Takeichi, 2014). In contrast, the SJs in invertebrate, or TJS in vertebrate, is mainly appreciated by their role in forming the paracellular diffusion (Harden et al., 2016; Tepass and Hartenstein, 1994). SJs contain ladder-like septa that span the intermembrane space (Lane and Swales, 1982; Tepass and Hartenstein, 1994). In *Drosophila* the SJs consist of a large multi-protein complex, the parts of which are commonly utilized in vertebrate TJs, locating apically to AJs (Harden et al., 2016; Izumi and Furuse, 2014).

Previous studies indicate that various *scrib* alleles show overproliferation phenotypes in the wing imaginal disc, but some alleles, e.g. *scrib^5^*, still appear to maintain ABP (Khoury and Bilder, 2020; Zeitler et al., 2004). The Scrib N-terminal leucine rich region (LRR) domain is necessary for both cell polarity and control of cell proliferation, while C-terminal PDZ domains enhance the ability of the LRR to localize SJs proteins and to provide full proliferation control (Fig. 1A) (Bonello and Peifer, 2019; Zeitler et al., 2004). However, the detailed molecular mechanisms behind this remain to be addressed.

**Fig. 1.**
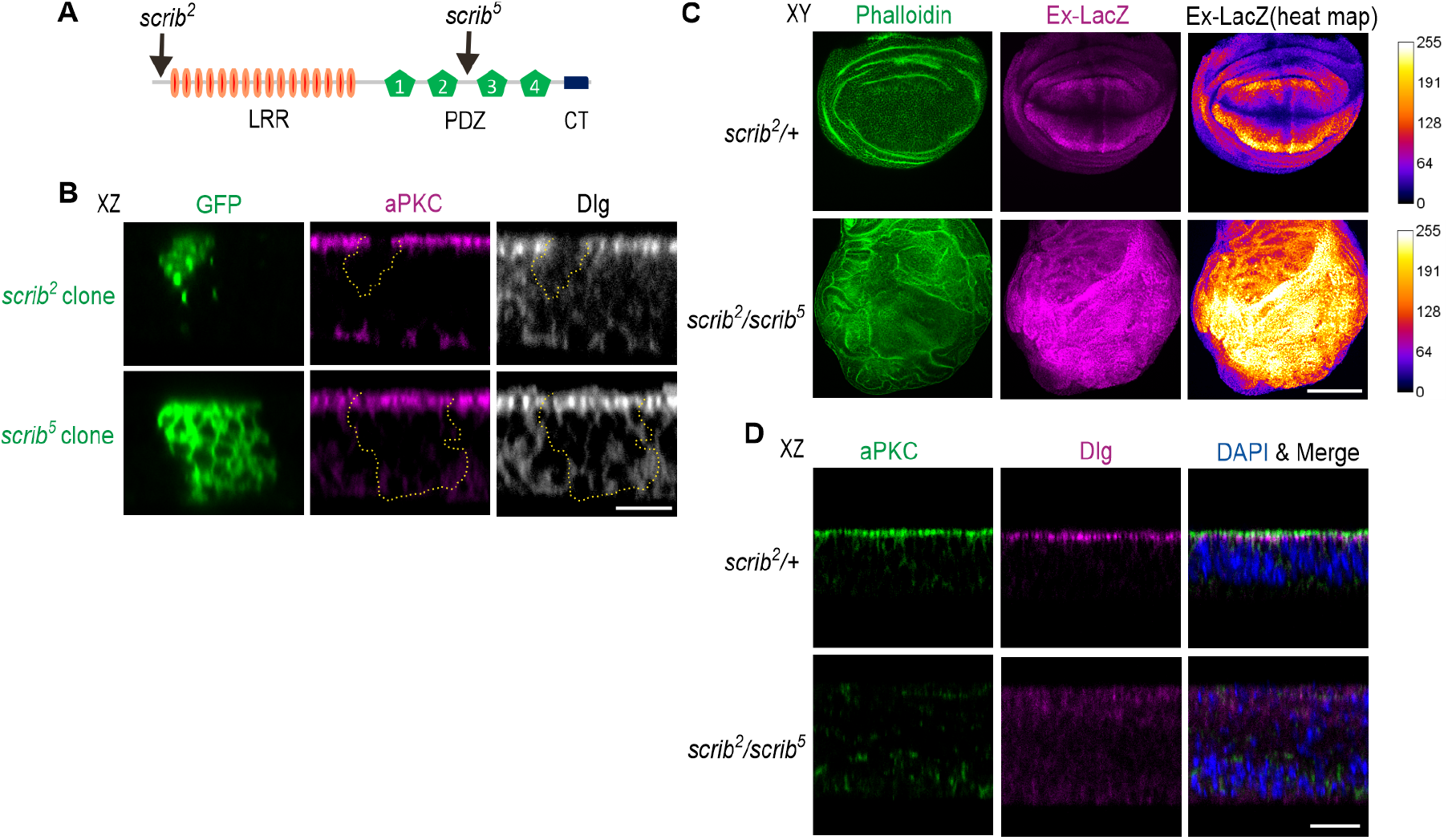
Scrib and apicobasal polarity in wing epithelial cells. (A) A schematic showing the secondary structure of Scrib, and the position where a premature stop codon is found in *scrib^2^* and *scrib^5^* mutations. Approximate positions of leucine rich region (LRR), Psd95-Dlg-Zo1 (PDZ) and C-terminal (CT) domains are labelled. (B) Representative transverse section (XZ) images showing mosaic studies of *scrib^2^* or *scrib^5^* mutant cells labelled with mCD8:GFP (green); the tissues were co-stained with aPKC (apical marker, magenta) and Dlg (basolateral marker, grey). Apical is up and basal bottom. (C) Representative coronal (XY) images showing the wing pouch of *scrib^2^/+* or *scrib^2^/scrib^5^* mutant animals, labelled with phalloidin (F-Actin, green) and *ex-lacZ* (Yki activity, magenta). Dorsal is up and ventral bottom. Anterior is left and posterior right. Heat map showing the intensities of LacZ staining. A scale for the heat maps is indicated on the right. The heat map was produced using the ROI color coder plugin, matching measurements to a color of a lookup table (LUT) of ImageJ/FIJI. (D) Representative transverse section (XZ) images showing the wing pouch of *scrib^2^/+* or *scrib^2^/scrib^5^* mutant animals, labelled with aPKC (green), Dlg (magenta) and merged with DAPI (nuclear marker, blue). Scale bars, 10 μm (B), 100 μm (C) and 20 μm (D).

In this study, to understand how ABP is regulated at the tissue level, we investigate Scrib in the wing imaginal disc and unveil an intercellular mechanism to coordinate ABP within a tissue. Our findings suggest that ABP coordination takes place either progressively or regressively through novel mechanism termed “intercellular alignment of ABP” in a context dependent manner.

## Results

### Scrib*^5^* contributes to apicobasal polarity in wing epithelial cells in a context-dependent manner

Previous studies indicate that one of the *scrib* allele, *scrib^5^*, shows overproliferation phenotypes in the wing imaginal disc but appears to maintain ABP, indicating that the N-terminal LRR domain might be sufficient for ABP, but not growth control (Khoury and Bilder, 2020; Zeitler et al., 2004). To investigate this further, we employed mosaic analysis with a repressible cell marker (MARCM), using two distinct *scrib* mutant alleles in the *Drosophila* wing imaginal disc (Fig. 1A) (Lee and Luo, 1999). When null mutant clones (*scrib^2^*) were introduced, only very small clone size was produced and accompanied by an autonomous reduction of expression of atypical protein kinase C (aPKC), an apical marker, and Dlg (Fig. 1B) (Chen et al., 2012; Kanda and Igaki, 2020). In contrast, clones of *scrib^5^*, which contains a stop codon between second and third PDZ domains, do not show significant changes of aPKC and Dlg spatial distribution (Fig. 1A, B). These results suggest that Scrib^5^ protein is sufficient for maintaining ABP in wing epithelial cells when *scrib^5^* cells are surrounded by wild type (WT) cells. In contrast, the wing discs from *scrib^5^* (*scrib^2^/ scrib^5^*) animals show overproliferation phenotypes accompanied by significant activation of Yki as shown by *ex-lacZ* expression that serves as a readout of Yki activity, along with increased apoptosis (Fig. 1C, D, Fig. S1A) (Pan et al., 2018). Our data also showed that distinct compartments of aPKC and Dlg dissipated (Fig. 1D), indicating that ABP is not properly maintained in *scrib^5^* tissues, and accordingly growth control appears to be defective. This suggests that *scrib^5^* cells *per se* are incompetent in maintaining ABP and primed for malignancy.

### Global regulation of apicobasal polarity in wing epithelial cells

Given that *scrib^5^* clone cells maintain ABP, we wondered whether ABP in *scrib^2^/ scrib^5^* tissues might be restored when full length *Scrib* (*Scrib^FL^*) is expressed in neighboring cells. To test this, Scrib^FL^ was expressed in a stripe of anterior cells abutting the anterior-posterior boundary (driven by *patched* (*ptc)-GAL4*) in the *scrib^2^/ scrib^5^* wing imaginal disc. Remarkably, tissue architecture was thoroughly restored, and Dlg distribution was largely normal across the tissue (Fig. 2A, B). Yki signal was also restored in the entire tissue (Fig. 2C). Similar results were observed when *scrib^FL^* was ectopically expressed in the dorsal compartment (driven by *apterous (ap)-GAL4*) of *scrib^2^*/ *scrib^5^* wing imaginal discs. Likewise, tissue architecture and Yki activity were largely rescued globally (Fig. S1B). Moreover, Dlg distribution and cell viability were also restored in both dorsal and ventral cells (Fig. S1A, C, D). These results suggest that cells expressing *scrib^FL^* have the capacity for ABP restoration in *scrib^2^/ scrib^5^* cells in a non-autonomous manner, and deregulation of growth control and tissue homeostasis are recovered as well. We therefore term this non-autonomous progression of cell polarity ‘intercellular alignment of apicobasal polarity (ABP)’.

**Fig. 2.**
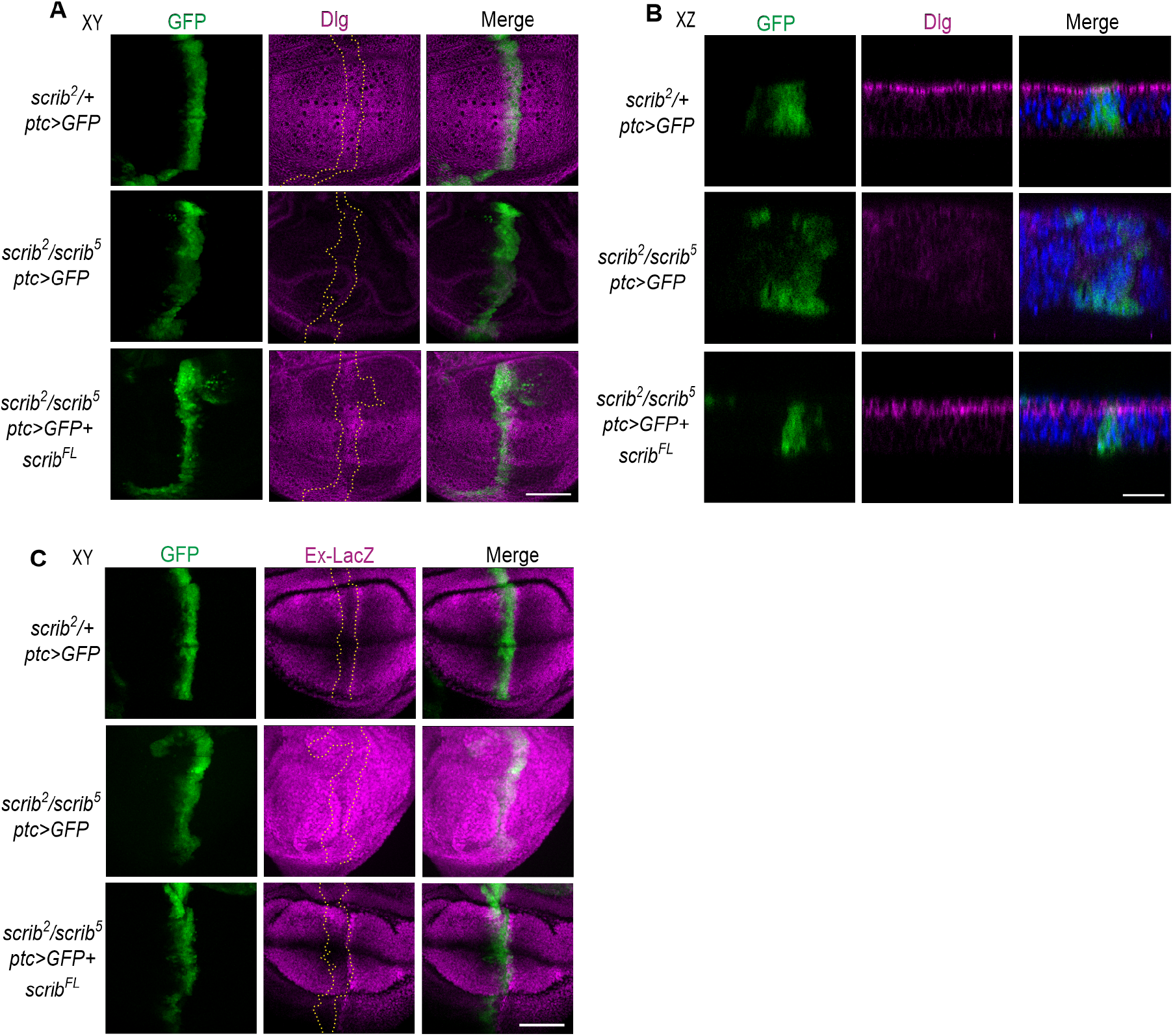
Oncogenicity of *scrib^5^* is restrained by neighboring WT cells. (A, B) Representative coronal (XY, E) or transverse section (XZ, F) images showing the wing pouch of *scrib^2^/+* or *scrib^2^/scrib^5^* mutant animals expressing GFP alone or GFP (green) and *scrib^FL^* driven by *ptc*-*GAL4* driver. Tissues were co-stained with Dlg (magenta) and DAPI (blue). Numbers of neoplastic wing imaginal discs have been seen in 0 out of 24 in *scrib^2^/+, ptc>GFP*; 28 out of 32 in *scrib^2^/scrib^5^, ptc>GFP*; 5 out of 34 in *scrib^2^/scrib^5^, ptc>GFP + scrib^FL^.* (C) Representative coronal (XY) images showing the wing pouch of *scrib^2^/+* or *scrib^2^/scrib^5^* mutant animals expressing GFP alone, or GFP and *scrib^FL^* driven by *ptc*-*GAL4* driver. Tissues were co-stained with *ex-lacZ* (magenta). The dashed lines outline GFP-positive cells (A, C). Scale bars, 50 μm (A, C), and 20 μm (B).

### Conditional *scrib* knockdown leads to non-autonomous loss of apicobasal polarity

Previous studies indicate that RNA interference (RNAi)-mediated knockdown (KD) of the Scrib module driven by *ptc-GAL4* sufficiently induced Yki dependent cell overproliferation and neoplasia in the wing imaginal disc (Chen et al., 2012; Ma et al., 2013). We wondered how ABP is affected prior to overproliferation upon *scrib* knockdown. To address this, we employed conditional knockdown of *scrib* by using temperature sensitive GAL80 (GAL80^ts^) in a combination with GAL4/UAS system (Fig. 3A). Strikingly, loss of Scrib was not only observed in *scrib* KD cells, but also in the anterior flanking cells 2 days (2D) after temperature shift (ATS) (Fig. 3B, C). The loss of Scrib further extends distally at 3D and 4D ATS (Fig. 3B-D). By 4D ATS, Scrib is massively depleted across the tissue, resulting in disorganization of tissue structure (Fig. 3B, C). Next, we addressed how tissue growth and homeostasis are regulated in such conditions. Our data reveal that initial reduction of Scrib involves the subsequent activation of Yki signal and increased apoptosis, followed by overproliferation that becomes evident at 4D ATS (Fig. 3B, C, E, Fig. S2). Since previous studies suggest that ectopic Yki activity is key for overproliferation after *scrib* KD (Yang et al., 2015), we investigated whether increased Yki activity is required for expansion of loss of Scrib in flanking cells. When Wts was co-expressed in *scrib* KD cells, increased *ex-lacZ* expression was curbed, and therefore, hyperplastic phenotypes were sufficiently suppressed (Fig. 3B-F). Under such conditions, loss of Scrib in the flanking cells remained detectable without further lateral deterioration (Fig. 3B-D), indicating that initial Scrib loss in the flanking cells occurs irrespective of Yki activity, then further expansion of Scrib loss is facilitated by increased Yki activity. Notably, with *wts* expression, *scrib*-KD cells remain integrated within the wing epithelia, suggesting that the global disorganization of tissue architecture can be largely attributed to local hyperactivation of Yki in ABP-compromised cells (Fig. 3B-F). Taken together, these data suggest that intercellular regressive alignment of ABP takes place from polarity-suboptimal cells to WT cells.

**Fig. 3.**
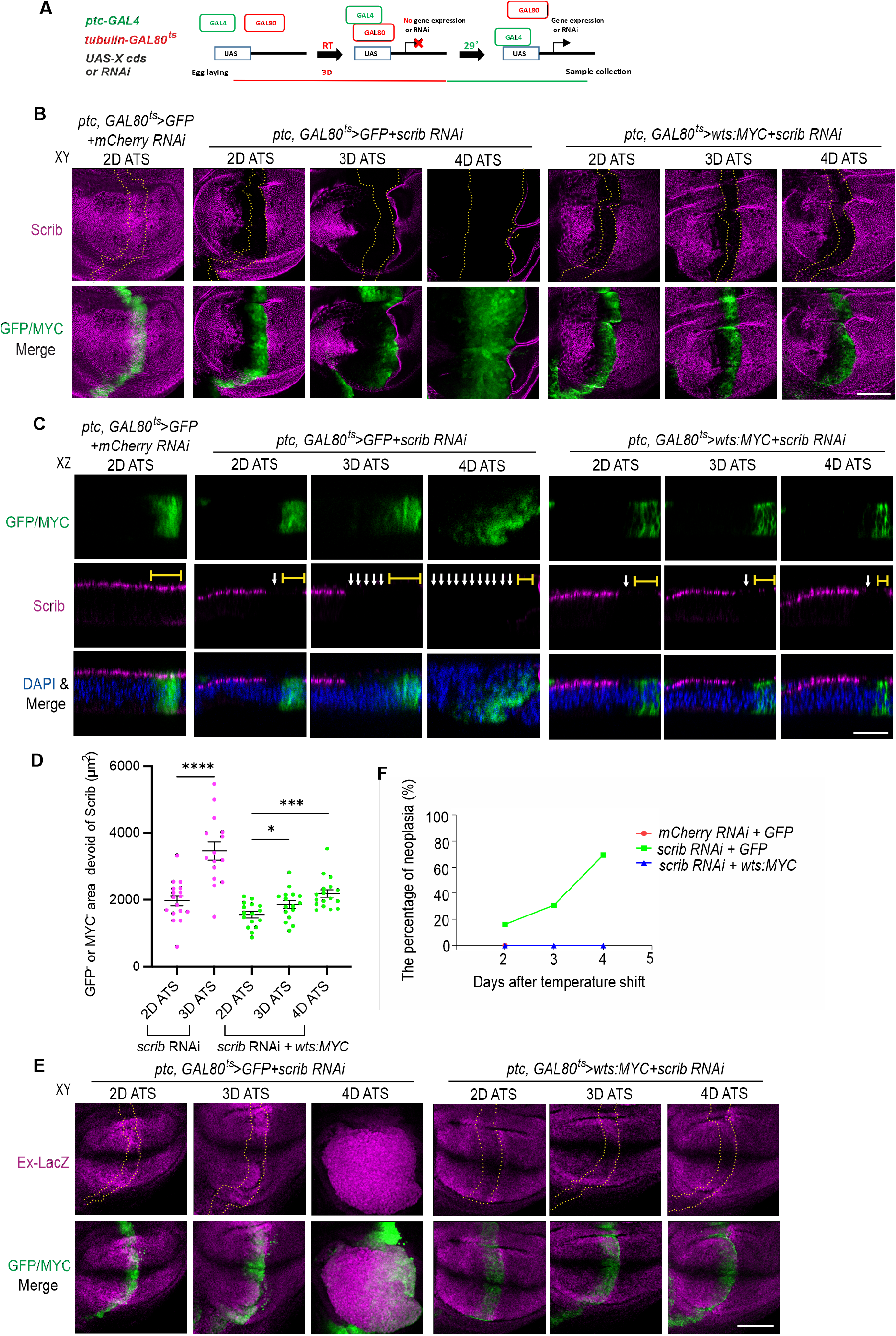
Non-autonomous loss of apicobasal polarity is instigated by conditional knockdown of *scrib*. (A) Schematic of conditional RNAi knockdown by using GAL80^ts^ together with the GAL4 system. (B, C) Representative coronal (XY, B) or transverse section (XZ, C) images showing the wing pouch with conditional expression of *GFP* and *mCherry RNAi, GFP* and *scrib RNAi*, or *wts:MYC* and *scrib RNAi* driven by *ptc-GAL4*. Tissues were collected between 2-4 days after temperature shift (ATS) for immunofluorescent analyses, labelled with Scrib (magenta), GFP or MYC (green), and DAPI (blue). The dash lines outline GFP/ MYC-positive cells (B). The brackets indicate the apical range of GFP/ MYC-positive cells and the arrows indicate loss of polarity (C). Please note that loss of Scrib is observed in the only anterior flanking region at 2D ATS and then in the posterior flanking region at 3D ATS or later when conditional *scrib RNAi* takes place. (D) Quantification of the area loss of Scrib outside GFP/ MYC positive cells in wing pouch. Sample sizes are 17 and 15 (*scrib* RNAi 2D and 3D ATS), and 16, 16 and 17 (*scrib* RNAi and *wts:MYC* 2D, 3D and 4D ATS). \**P* = 0.0462, ***P = 0.0002, \*\*\*\**P* < 0.0001. Data are means ± 95 % confidence intervals (CIs). Statistical significance was calculated by the two-tailed *t*-test. (E) Representative coronal (XY) images showing the wing pouch with conditional expression of *GFP* and *scrib RNAi*, or *wts:MYC* and *scrib RNAi* driven by *ptc*-*GAL4*. Tissues were collected between 2-4 days after temperature shift (ATS) for immunofluorescent analyses, labelled with *ex-lacZ* (magenta), and GFP or MYC (Green). (F) Quantification of the percentage of the neoplasia. Sample sizes are 48 (*mCherry* RNAi 2D ATS), 74, 78 and 68 (*scrib* RNAi 2D, 3D and 4D ATS), and 54, 58 and 54 (*scrib* RNAi and *wts:MYC* 2D, 3D and 4D ATS). Scale bars, 50 μm (B, E), and 30 μm (C).

### α-Catenin genetically and physically interacts with Scrib

Next, we address the molecular mechanisms underlying ABP regression through intercellular alignment. Cell-cell communication of epithelial cells is often mediated through AJs (Pinheiro and Bellaïche, 2018). Therefore, we first tested how key components of AJs are affected after conditional *scrib* KD (2D ATS). DE-Cadherin (DE-Cad) and β-Catenin (β-Cat) remain intact, although their expression is significantly decreased after *DE-cad* KD (Fig. 4A, B, Fig. S3A). In contrast, α-Catenin (α-Cat) expression significantly decreased in either *scrib* or *DE-cad* KD (Fig. 4A, B). α-Cat was synergistically depleted when *scrib* and *DE-cad* were simultaneously knocked down (dKD; Fig. 4A, B). We also confirmed that α-Cat biochemically interacts with LRR domains and PDZ3/4 domains of Scrib (Fig. S3B, C). These results suggest that α-Cat may be a thus far insufficiently characterized factor involved in the regulation of the Scrib complex. In contrast, conditional KD of α-Cat reveals that Scrib distribution was barely affected by 2D ATS (Fig. 4C), and could not be investigated further due to cell death and tissue malformation from 3D ATS onwards. Given that Scrib complex is enriched around and functions as co-factor of SJs (Harden et al., 2016; Rice et al., 2021), we wondered whether core-components of SJs, Scrib complex and α-Cat mutually interact to sustain intercellular alignment of ABP. To corroborate this, we examined how Scrib distribution is affected by depleting α-Cat and SJs components. Conditional double KD (dKD) of *α-Cat* and SJs components (*NrxIV* or *Kune*) driven by *ptc-GAL4* indeed leads to reduced expression of Scrib only in a part of cells abutting the anterior-posterior boundary by 2D ATS (Fig. 4C). Conditional dKD beyond 2D ATS again results in tissue malformation, thus it was difficult to conclude whether reduced Scrib expression in dKD cells further lead to non-autonomous Scrib reduction or intercellular progressive alignment of ABP from abutting WT cells is more dominant than intercellular regressive alignment of ABP in KD cells. To address this, we employed *nubbin (nub)-GAL4* whose activity is across the whole wing pouch to perform the KD experiments. Interestingly, *α-Cat* KD gave rise to patchy loss of Scrib while most cells seem unaffected (Fig. S3D). However, upon dKD, a vast majority of cells within the wing pouch were devoid of Scrib (Fig. S3D). These data suggest that α-Cat can genetically interact with SJs components to cell-autonomously regulate Scrib localization, and the preserved ABP in GFP-positive cells of *ptc*-*GAL4*-mediated dKD is very likely due to the instructive cue derived from WT cells through intercellular progressive alignment of ABP. Altogether, these results clearly indicate the genetic interactions between α-Cat, Scrib and core-components of SJs.

**Fig. 4.**
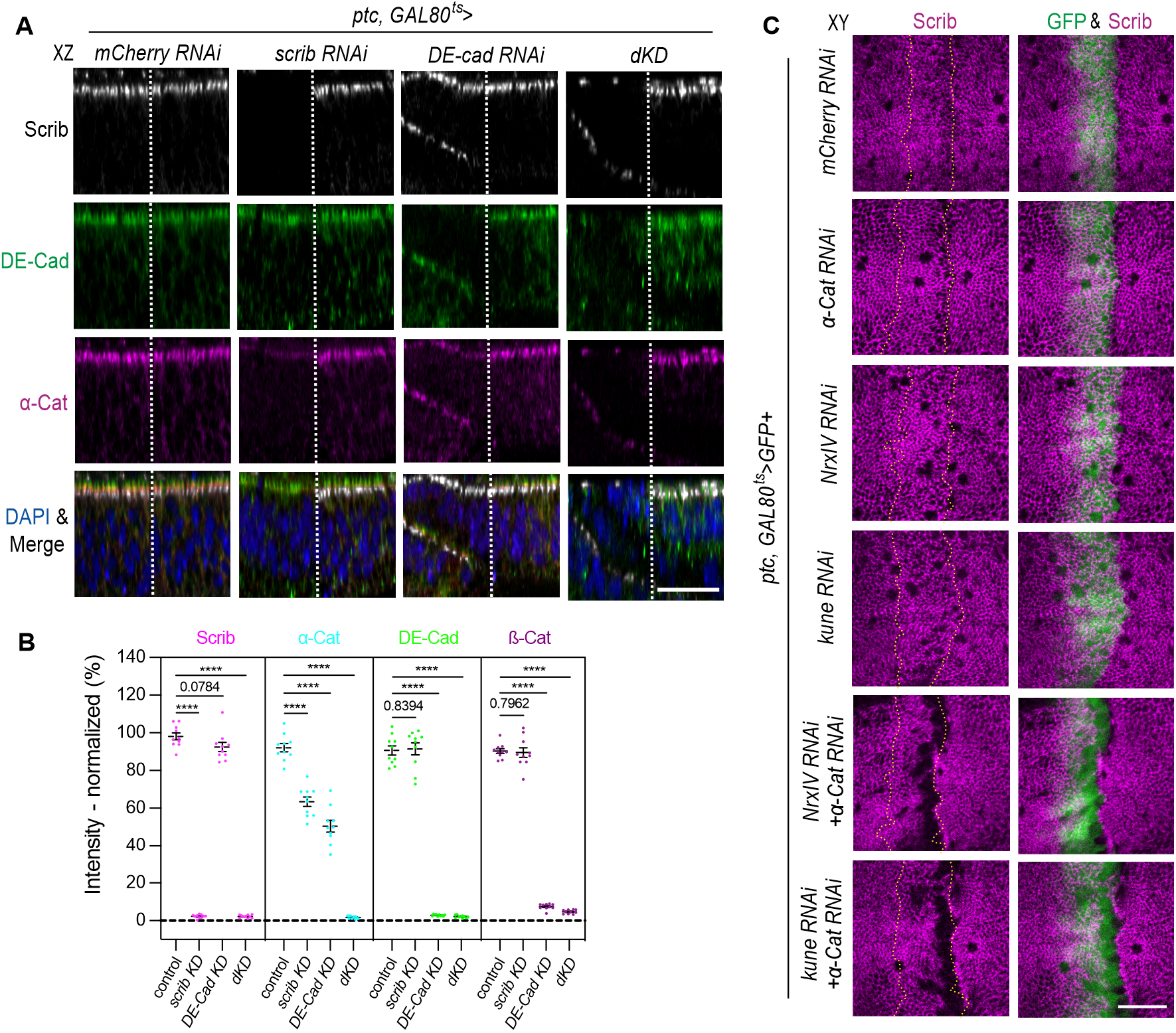
α-Catenin associates with Scrib. (A) Representative transverse section (XZ) images showing the wings imaginal disc with conditional expression of *mCherry* RNAi alone (control), *scrib* RNAi (*scrib* KD), *DE-cad RNAi* (*DE-cad* KD) or *scrib* / *DE-cad RNAi* (dKD) driven by *ptc-GAL4*. Tissues were collected two days after temperature shift (ATS) for immunofluorescent analyses, labelled with Scrib (grey), DE-Cad (green), α-Cat (magenta) and DAPI (blue). The dash lines outline the anterior-posterior boundary. Anterior is to the left. Anterior domain in the figure corresponds to *ptc* expression. (B) Fluorescent intensity measurements of Scrib, α-Cat, DE-Cad and ß-Cat in the *ptc* expression area with RNAi, normalized to that of the abutting posterior area. Sample sizes are 10 in each quantification. \*\*\*\**P* < 0.0001. Data are means ± 95 % confidence intervals (CIs). Statistical significance was calculated by the two-tailed *t*-test. (C) Representative coronal (XY) images showing the wings with conditional expression of *mCherry RNAi* alone, *α-Cat RNAi* and/ or *NrxⅣ RNAi, α-Cat RNAi* and/ or *kune RNAi* driven by *ptc-GAL4*. Tissues were collected at 2 days after temperature shift (ATS) for immunofluorescent analyses, labelled with Scrib (magenta) and GFP (green). The dash lines outline GFP-positive cells. Scale bars, 20 μm (A, C).

### Association of α-Catenin with Scrib and intercellular alignment of apicobasal polarity

How is α-Cat involved in Scrib signaling? Previous studies suggest that Scrib-LRR domains are indispensable for ABP establishment, while Scrib-PDZ domains support it by stabilizing cortical Scrib (Zeitler et al., 2004). We therefore hypothesize that α-Cat may help stabilize Scrib localization at the cortical domain through interaction with PDZ domains. To test this hypothesis, we generated and expressed various constructs in the wing imaginal disc. When various Scrib proteins (MYC-tagged C-terminally) were overexpressed under the control of *ptc-GAL4*, we observed that full-length Scrib (Scrib^FL^:MYC) becomes more dominantly localized at the cortical domain where GFP trapped Scrib (Scrib:GFP) is located, while LRR domains alone (Scrib^LRR^:MYC) affects Scrib:GFP but also shows broader distribution at the cell periphery (Fig. S3E, F). Interestingly, LRR:α-Cat chimera (Scrib^LRR^:α-Cat:MYC) recapitulates the pattern of Scrib^FL^ and outcompetes Scrib:GFP (Fig. S3E, F). Nevertheless, α-Cat alone (α-Cat:MYC) or Scrib variant lacking LRR domain (Scrib^dLRR^:MYC) is expressed rather broadly and barely affects Scrib:GFP (Fig. S3E, F). Taken together these data suggest that the association of α-Cat facilitates cortical localization of Scrib. Next, we hypothesized that α-Cat localizing around SJs, but not AJs, may play a role in sustaining Scrib function. To test this, we generated multiple constructs to compensate loss of *scrib*, including chimeras containing either Kune or DE-Cad, with α-Cat (Kune:α-Cat or DE-Cad:α-Cat). Overproliferation phenotypes caused by *scrib* KD were sufficiently restored by Scrib^FL^ expression as expected (Fig. 5A). Although restoration was not observed when either α-Cat or Kune was expressed, Kune:α-Cat chimera protein significantly rescued tissue malformation and ectopic Yki activity within the wing pouch (Fig. 5A-C). Furthermore, loss of Scrib in the flanking cells from the conditional *scrib* KD is still visible 2D ATS, however, successive loss of Scrib was significantly suppressed by co-expression of the Kune:α-Cat chimera (Fig. 5B, D). Importantly, such restoration phenotypes were not observed when DE-Cad:α-Cat chimera was expressed (Fig. 5A-C), highlighting the importance of subcellular localization of α-Cat to SJs during intercellular alignment of ABP.

**Fig. 5.**
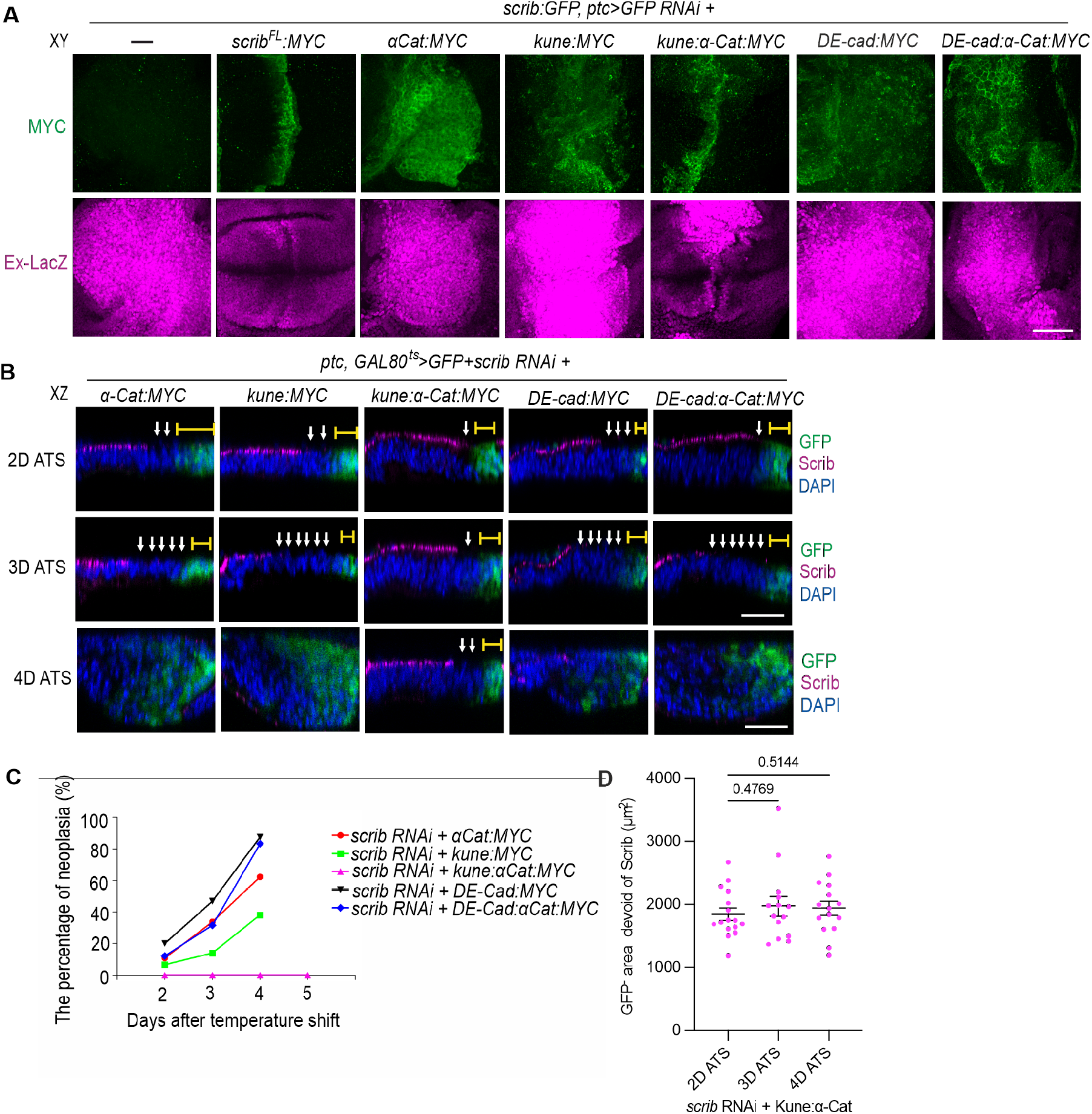
α-Catenin associates with Scrib to sustain intercellular alignment of apicobasal polarity. (A) Representative coronal (XY) images showing the wing pouch with expression of *GFP RNAi* and *Scrib^FL^*, *α-Cat*, *kune*, *kune:α-Cat*, *DE-cad*, or *DE-cad:α-Cat* driven by *ptc*-*GAL4* in GFP-trapped animals (*scrib:GFP*). Tissues were collected for immunofluorescent analyses, labelled with MYC (green) and Ex-lacZ (magenta). (B) Representative transverse section (XZ) images showing the wings with conditional expression of *GFP* and *scrib RNAi* together with *α-Cat, kune, kune:α-Cat, DE-cad* or *DE-cad:α-Cat* driven by *ptc-GAL4*. Tissues were collected at 2-4 days after temperature shift (ATS) for immunofluorescent analyses, labelled with Scrib (magenta), GFP (green) and DAPI (blue). The brackets indicate the apical range of GFP-positive cells and the arrows indicate loss of polarity. (C) Quantification of the percentage of the neoplasia. Sample sizes are 74, 71 and 61 (*scrib* RNAi and *α-Cat:MYC* 2D, 3D and 4D ATS), 62, 43 and 55 (*scrib* RNAi and *kune:MYC* 2D, 3D and 4D ATS), 63, 54, 79 and 67 (*scrib* RNAi and *kune:α-Cat:MYC* 2D, 3D 4D and 5D ATS), 40, 64 and 64 (*scrib* RNAi and *DE-Cad:MYC* 2D, 3D and 4D ATS), and 58, 38 and 54 (*scrib* RNAi and *DE-Cad:α-Cat:MYC* 2D, 3D and 4D ATS). (D) Quantification of the area loss of Scrib outside GFP positive cells in wing pouch. Sample sizes are 16, 14 and 15 (*scrib* RNAi and *kune:α-Cat* 2D, 3D and 4D ATS). Data are means ± 95 % confidence intervals (CIs). Statistical significance was calculated by the two-tailed *t*-test. Scale bars, 50 μm (A) and 30 μm (B).

### The dosage of Scrib affects horizontal communication between optimal and sub-optimal cells

Our data reveal that intercellular regressive alignment of ABP is regulated by mutual communication between cells, and ABP regression in WT cells occurs in a graded manner, indicating the quantity of Scrib complex may be crucial. To understand this, we knocked down *scrib* in flies expressing *scrib* at various levels. When these experiments were tested in flies with one null allele of *scrib* (*scrib*^2^/+), which develops normally, ABP loss in surrounding cells (*scrib*^2^/+) accelerates and propagates throughout the wing pouch by 3D ATS (Fig. 6A-C). Accordingly, tissue disorganization and neoplasia were frequently observed by 3D ATS (Fig. 6A, B, D). The haploinsufficiency of *scrib* that is not detectable during homeostasis is manifested when *scrib*^2^/+ cells are facing the ABP-compromised cells, suggesting both amount and localization of Scrib protein is important to maintain robustness of ABP. To validate this further, we used tissue with ectopic Scrib by expressing exogenous Scrib under the control of the polyubiquitin promoter (*ubi-scrib*). Interestingly, even though ABP loss in the cells abutting *scrib* KD cells still occurs by 2D ATS, no further progression was detected even by 5D ATS (Fig. 6A-D). These results reveal that the local loss of function of *scrib* differentially impact on tissue architecture and overall the growth within the tissue with different dosages of *scrib*.

**Fig. 6.**
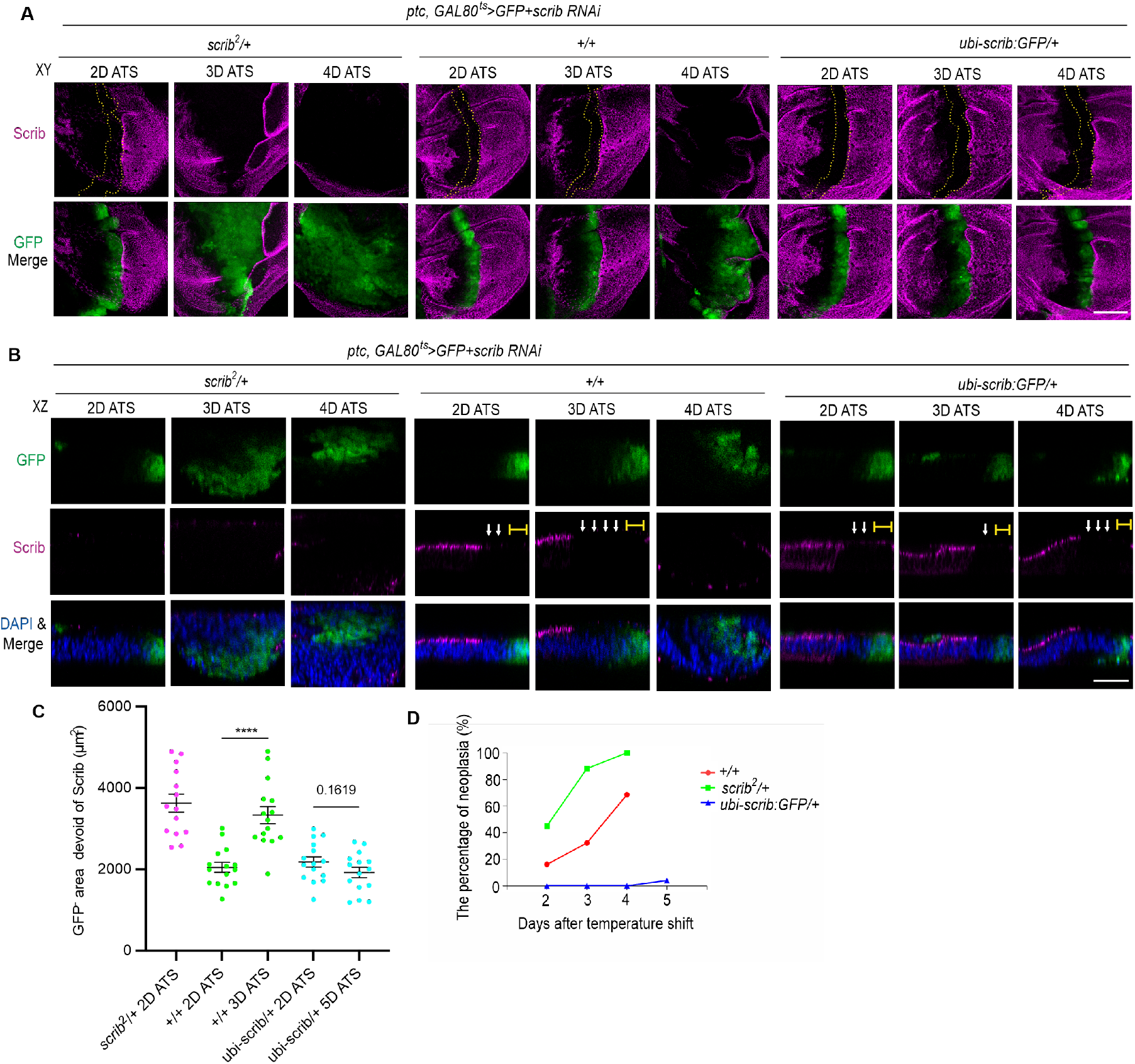
Non-autonomous loss of apicobasal polarity depends on the dosage of Scrib. (A, B) Representative coronal (XY, A) or transverse section (XZ, B) images showing the wing pouch with conditional expression of *GFP* and *scrib RNAi* driven by *ptc*-*GAL4* in *scrib^2^/+*, +/+ or *ubi-scrib/+* animals. Tissues were collected between 2-4 days after temperature shift (ATS) for immunofluorescent analyses, labelled with Scrib (magenta), GFP (green) and DAPI (blue). The dash lines outline GFP-positive cells (A). The brackets indicate the apical range of GFP-positive cells, and the arrows indicate loss of polarity (B). (C) Quantification of the area loss of Scrib outside GFP positive cells in wing pouch. Sample sizes are 14 (*scrib^2^*/+ 2D ATS), 15 and 15 (+/+ 2D and 3D ATS), and 15 and 15 (*ubi-scrib*/+ 2D and 5D ATS). \*\*\*\**P* < 0.0001. Data are means ± 95 % confidence intervals (CIs). Statistical significance was calculated by the two-tailed *t*-test. (D) Quantification of the percentage of the neoplasia. Sample sizes are 56, 62 and 54 (*scrib* RNAi in +/+ 2D, 3D and 4D ATS), 40, 85 and 63 (*scrib* RNAi in *scrib^2^*/+ 2D, 3D and 4D ATS), and 42, 35, 38 and 50 (*scrib* RNAi in *ubi-scrib/+* 2D, 3D 4D and 5D ATS). Scale bars, 50 μm (A), 30 μm (B).

## Discussion

Here, our data reveal that a novel mechanism named “intercellular alignment of ABP” may play a crucial role in regulating epithelial homeostasis and undermining epithelial integrity. Importantly, intercellular alignment of ABP takes place either progressively or regressively in a context-dependent manner. Although ABP maintenance has been recognized as one of the means for epithelial homeostasis, molecular mechanisms underlying ABP have been largely focused on the intracellular compartment. Our findings open the horizon of intercellular mechanisms of ABP through horizontal communications among optimal and sub-optimal cells. Cellular communication between optimal and sub-optimal cells has been previously investigated, demonstrating that cell competition plays a crucial role in tissue homeostasis (Baker, 2020; Bowling et al., 2019; Kanda and Igaki, 2020). In contrast to the competition paradigm, in which sub-optimal/loser cells are eliminated to maintain tissue homeostasis or restrain hyperplasia, our studies show that intercellular alignment of ABP serves as a novel means that coordinates integration of cells through intercellular ABP regulation.

Our studies employing conditional RNAi KD show that ABP in WT cells can be instigated when they are adjacent to *scrib*-KD cells through intercellular regressive alignment of ABP (Fig. 7A). Remarkably, this process is significantly affected by dosage of Scrib, further suggesting the active engagement of intercellular alignment of ABP in physiological and pathological conditions. Our data also reveal that α-Cat plays a key role in sustaining intercellular regulation of ABP. In contrast to previously proposed α-Cat localization to AJs (Takeichi, 2014), our findings reveal that α-Cat appears to be localized at SJs for ABP regulation as follows. First, cellular localization of α-Cat is partly regulated by Scrib. Second, α-Cat biochemically interacts with Scrib. Third, SJs-but not AJs-localized α-Cat partially rescue loss of *scrib* function. Since α-Cat, as a scaffold protein, displays multiple interactions and functions together with other proteins, loss of function of *α-cat* does not show clear cut phenotypes (Takeichi, 2014). Here, our data sufficiently support that α-Cat is involved in intercellular alignment of ABP.

**Fig. 7.**
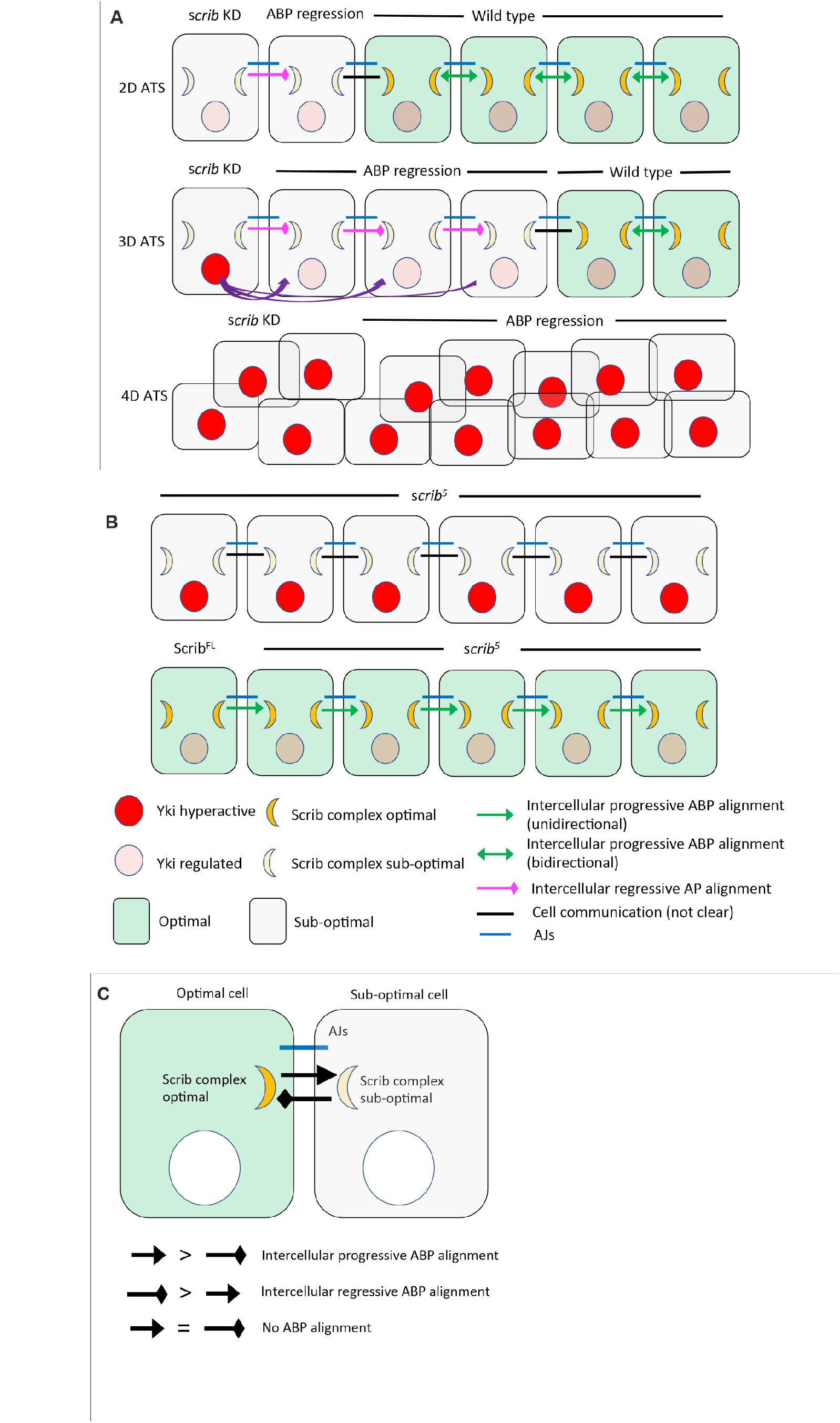
Schematics of intercellular apicobasal polarity alignment. (A) Schematics of intercellular regressive ABP alignment. *scrib* KD cells lead to loss of ABP in flanking wild type cells (upper panel). Yki signal after *scrib* KD further facilitates loss of ABP distally (middle panel). Eventually most cells become neoplastic, with hyperactivation of Yki (bottom). (B) Schematics of intercellular progressive ABP alignment. Partial loss of function of *scrib* complex, e.g. *scrib^5^*, is intrinsically premalignant. Cells expressing Scrib^FL^ restore ABP in suboptimal cells in a non-autonomous manner to maintain tissue homeostasis, while suboptimal cells contain Scrib^5^ protein that serves as a “seed” for restoring ABP. (C) Cell-cell communication between optimal and ABP sub-optimal cells. When progressive ABP alignment is more dominant than regressive one, intercellular progressive ABP alignment takes place, and vice versa.

Since expression of ABP determinants can be targeted by both pathogenic assaults and normal cellular programs, e.g. cell division and mesenchymal-epithelial transition (EMT) (Bonello and Peifer, 2019; Javier, 2008; Javier and Rice, 2011; Varga and Greten, 2017; Yamben et al., 2013), intercellular alignment of ABP appears to be important for tissue integrity. Several factors including those regulating cell viability and proliferation may bias one side in this ‘tug of war’ between optimal (wild type) cells and suboptimal *scrib*-KD cells. Besides the proper distribution of ABP determinants, their amount may be important for optimal cells to defend against premalignant cells’ loss of ABP. These factors can be key to determine the competitiveness of cells in instructing neighbors during intercellular alignment of ABP, safeguarding ABP in wild type cells and tissue architecture, but facilitating clonal expansion and drift of malignant cells and neoplasia.

In the case of *scrib^5^* mutants, tissue homeostasis is safeguarded in a different mechanism from cell competition. We postulate that intercellular progressive alignment of ABP requires “seeds” in recipient cells, since our data reveal that a mildly truncated form of Scrib appears to be needed for ABP alignment (Fig. 7B). In contrast, *scrib^2^* cells cannot establish ABP due to their lacking minimal Scrib protein elements, most likely LRR domain of Scrib, and are subject to elimination through cell competition (Chen et al., 2012; Kanda and Igaki, 2020). Therefore, cells with defective ABP but containing the seed element can still maintain ABP by receiving instructive cues. Although precise molecular mechanisms behind intercellular alignment of ABP remain to be addressed, we presume that intercellular alignment of ABP takes place through direct cell-cell contact, therefore either progression or regression of ABP is forwarded by relaying nodes. When progressive cues are more dominant than regressive ones, intercellular progressive ABP alignment takes place, and vice versa (Fig. 7C), revealing that novel type of competition between optimal and sub-optimal cells. One potential cellular target for intercellular alignment of ABP is SJs. Further studies will unveil more details about intercellular alignment of ABP.

Given that the maintenance of ABP across the tissue is a highly dynamic process, and epithelial cells require frequent remodeling of ABP during some physiological processes, including cell division and migration, the mechanism we unveiled in this work is essential to preserve robust ABP and thus epithelial integrity. However, it can be detrimental to the tissue when hijacked by malignant cells. The prospective efforts in investigating the underlying molecular underpinnings are critical to better understand global ABP maintenance and its relevance in diseases.

## Materials and Methods

### Fly genetics and husbandry

UAS-*mCherry* RNAi (#35785) UAS-*wts:MYC (#44250)*, *ex-lacZ (#44248)*, *nub*-*GAL4* (#25754), *apterous*-*GAL4* (#3041), UAS-*scrib* RNAi (#35748), *scrib^2^* (#41775), UAS-*NrxIV* RNAi (#39071), UAS-*GFP* RNAi (#44415) and *tub*P-*GAL80^ts^* (#7017) were obtained from the Bloomington Drosophila Stock Center (BDSC). UAS-*kune* RNAi (v108224), UAS-*shg/DE-cad* RNAi (v8024) were obtained from the Vienna Drosophila Resource Center (VDRC). *Scrib:GFP* (CA07683) was obtained from Fly Trap projects. FRT^82B^-*scrib^5^* was from D. Bilder (Zeitler et al., 2004), and *ex-lacZ, ptc*-*GAL4*, UAS-*EGFP* was from G. Halder (Yang et al., 2015).

Fly stocks were raised at 25°C unless otherwise mentioned. To induce MARCM clones, larvae were heat-shocked for two hours at 37°C at 72 hours after egg laying (AEL), followed by wing imaginal disc collection two days after heat shock.

Larvae for conditional knockdown mediated by *ptc-GAL4* and *nub-GAL4* were raised at 21℃ for 3 days before transferred to 29℃ and 27℃, respectively, followed by wing imaginal disc collection on the date indicated. Calculations for developmental timing at different temperatures were based on previously published data (Ashburner et al., 2004).

### Antibodies, chemicals and immunohistochemistry

Wing imaginal disc dissection and fixation were conducted as described previously (Gui et al., 2016). The following primary antibodies were used: rabbit anti-Scrib (1:200) from C. Doe (Albertson and Doe, 2003), mouse anti-Dlg (4F3, 1:50), rat anti-DE-Cad (DCAD2, 1:50), rat anti-α-Cat (DCAT-1, 1:50) and mouse anti-β-Cat (N2 7A1, 1:50) from Developmental Studies Hybridoma Bank (DSHB), rabbit anti-MYC tag (sc-789,1:1000 for WB and 1:100 for IF) and rabbit anti-aPKC (sc-937, 1:100) from Santa Cruz Biotechnology, mouse anti-β-Galactosidase (Z378A, 1:500) from Promega, mouse anti-MYC tag (#2276, 1:200 for IF), rabbit anti-cleaved-Dcp-1 (#9578, 1:200) from Cell Signaling Technology, mouse anti-GFP (MAB3580, 1: 5000) from Millipore, and mouse anti-γ-tubulin (T5326, 1:5000) from Sigma-Aldrich.

Secondary antibodies (1:200) and chemicals were as follows: goat anti-mouse IgG Alexa 488 (A11001), anti-mouse IgG Alexa 568 (A11004), goat anti-mouse IgG Alexa 647 (A21236), anti-rabbit IgG Alexa 568 (A11011), anti-rabbit IgG Alexa 647 (A21244) goat anti-rat IgG Alexa 488 (A11006) and Alexa 488 conjugated phalloidin (A12379) were purchased from Thermo Fisher Scientific, and DAPI (D9542) from Sigma-Aldrich.

### DNA constructs

All coding sequences (CDS) were amplified through PCR, tagged with MYC or GFP at the C-terminus and cloned into the *pUASt.attB (Bischof et al., 2007)*. The constructs encoding the Scrib fragments were generated previously (Gui et al., 2016). α-Cat was cloned into BglII and KpnI, Kune into KpnI and XbaI, and DE-Cad into NotI and KpnI sites of *pUASt.attB*, respectively. The chimeric constructs were generated by inserting α-Cat CDS into the C-terminus of Scrib^LRR^, Kune or DE-Cad (lacking the cytosolic domain, aa1-1348) before the stop codon.

### Transgenic flies

The plasmids for transgenesis were constructed as described above. Transgenic fly stocks were generated by injecting the plasmids into *yw*; (PBac y[+]-attP VK00018) embryos by BestGene Inc.

### Cell culture and production of recombinant proteins

*Drosophila* S2 cells were used to produce recombinant proteins for Western blotting (WB) and immunoprecipitation (IP) as previously described(Gui et al., 2016). The plasmids containing indicated CDS and *tub*P-*GAL4* were co-transfected into the cells using FuGENE HD transfection reagent (E2311, Promega) according to the manufacturer’s protocol, followed by cell harvest and lysis in IP lysis buffer (25mM Tris-HCl pH 7.4, 150 mM NaCl, 1% NP-40, 1 mM EDTA, 5% glycerol) 3 days after transfection. Cell lysates were subjected to WB and IP using the GFP-Trap^®^ Agarose (sta-100, ChromoTek) following the manufacturer’s protocol. All biochemical data shown are representative of three independent assays.

### Imaging and image analysis

Fluorescent images were obtained with a Zeiss LSM700 upright or Leica SP8 upright confocal microscope. All images were processed and analysed using ImageJ/FIJI (NIH, https://imagej.nih.gov/ij/). The images presented were composites of stacks (maximum intensity projection) unless otherwise specified. Only linear methods were applied. To measure the Scrib-negative area outside the GFP or MYC-positive domain in the wing pouch, the region was manually outlined and measured using the selection tools in ImageJ/FIJI. For the measurement of relative intensity of Scrib, DE-Cad, α-Cat, β-Cat and Scrib:GFP, a 20 μm x 100 μm area of *ptc*-*GAL4* apical domain (∼6 micron-thickness) was selected and subjected to intensity calculation using ImageJ/FIJI. Obtained values were subsequently standardized to that of the abutting posterior region.

### Statistics

Statistical analyses were performed using GraphPad Prism software (v.9.0.2, GraphPad). The number for all quantified data is indicated in the figure legends. Data are means ± 95 % confidence intervals (CIs). Statistical significance was calculated by the two tailed *t*-test method.

## Acknowledgments

We are grateful to Martti Montanari, Tamsin Samuels and Tambet Tõnissoo for thoughtful comments on the manuscript. We thank the Light Microscopy Unit of the Institute of Biotechnology of the University of Helsinki for their support. We thank D. Bilder, G. Halder, and C. Doe for fly stocks and antibodies. This work was supported by grant 308045 from the Academy of Finland, the Sigrid Juselius Foundation to O.S., and the Center of Excellence in Experimental and Computational Developmental Biology from the Academy of Finland to O.S. and M.M.

## Author Contributions

J.G., Y.H. and O.S. conceived the project and planned experiments. J.G. and Y.H. performed experiments. J.G., Y.H. and O.S. analyzed the results. S-M.M. and M.M. provided inputs. J.G., Y.H. and O.S. wrote the manuscript and all authors made comments. O.S. supervised the project.

## Supplementary figures

**Fig. S1.**
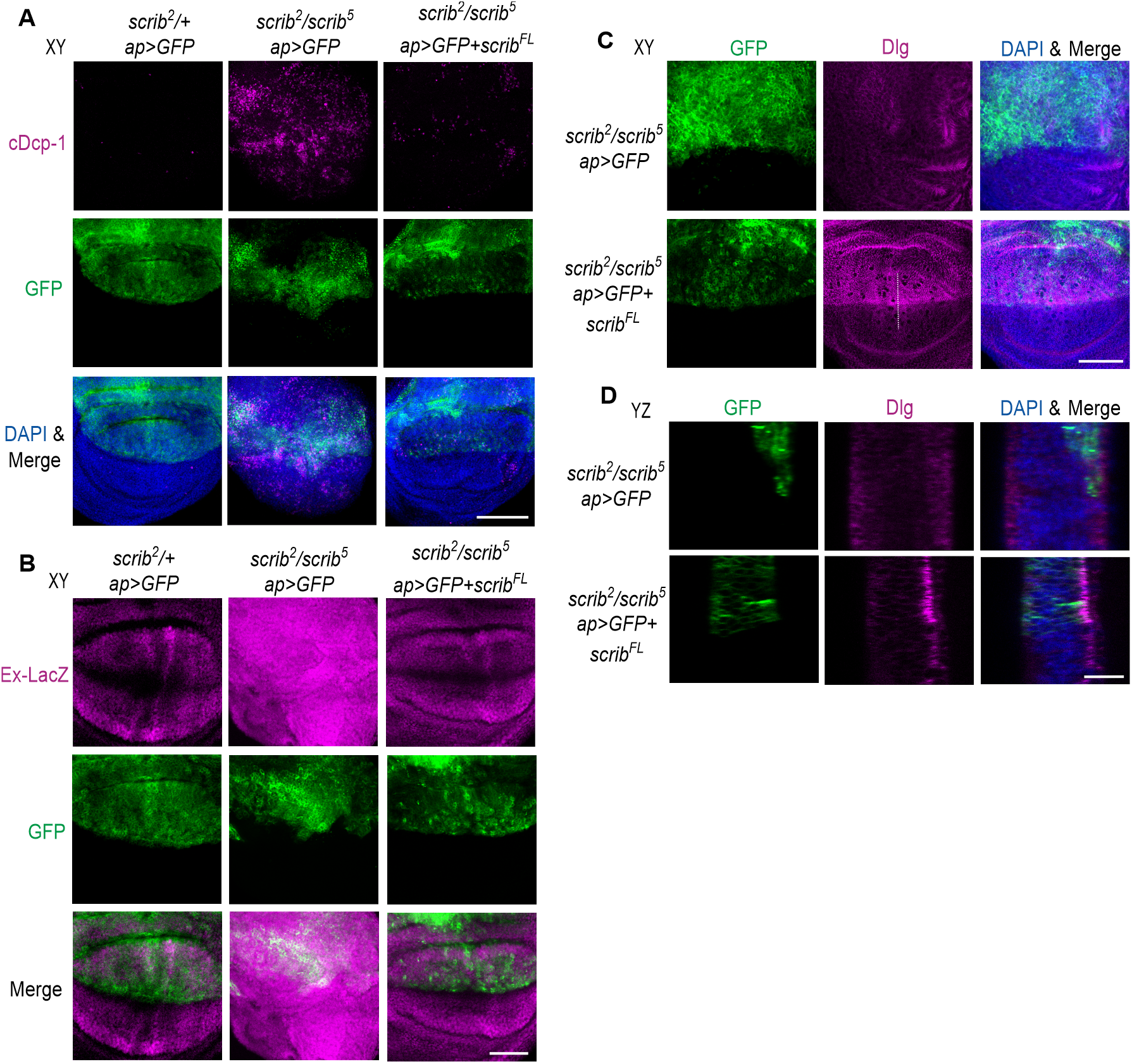
(A-D) Representative coronal (XY, A-C) or sagittal section (YZ, D) images showing the wings expressing *GFP* with or without *scrib^FL^* driven by *ap*-GAL4 in the background of *scrib^2^/+* or *scrib^2^/scrib^5^*, labelled with GFP, cDcp-1 (apoptosis marker, A), *ex-lacZ* (Yki activity, B), Dlg (basolateral marker, C, D) and DAPI (A, C, D). The dash lines (C) indicate the source of YZ sections (D). Numbers of neoplastic wing imaginal discs have been seen in 0 out of 26 in *scrib^2^/+, ap>GFP*; 28 out of 31 in *scrib^2^/scrib^5^, ap>GFP*; 1 out of 19 in *scrib^2^/scrib^5^, ap>GFP + scrib^FL^.* Dorsal is up and ventral bottom, anterior is left and posterior right (A-D). Scale bars, 100 μm (A), 50 μm (B, C), 20 μm (D).

**Fig. S2.**
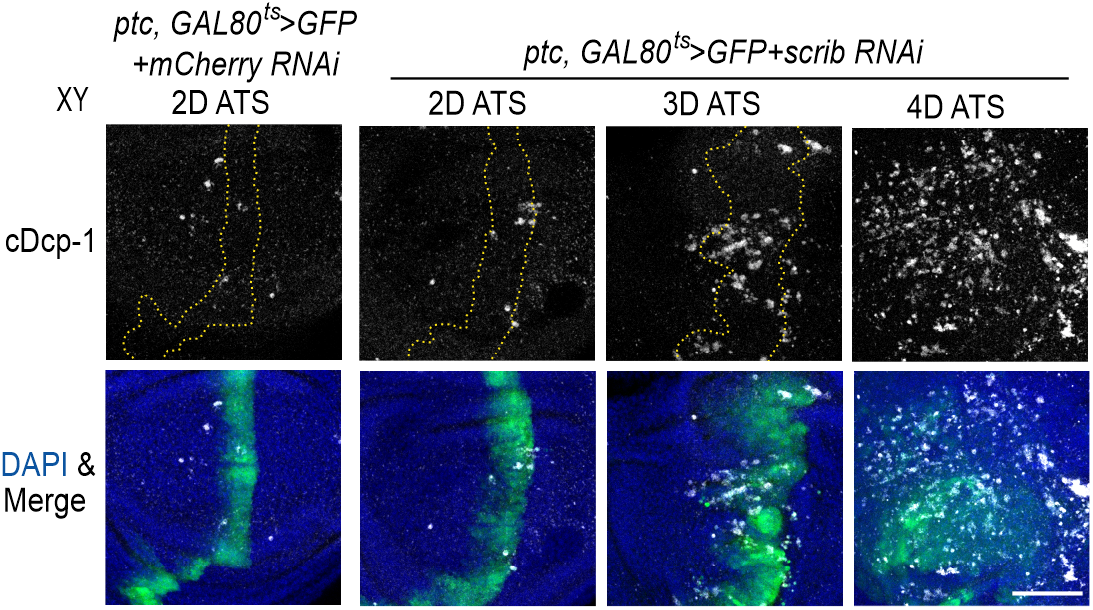
Representative coronal (XY) images showing the wing pouch with conditional expression of *GFP* and *mCherry RNAi* or *scrib RNAi* driven by *ptc*-*GAL4*. Tissues were collected between 2-4 days after temperature shift (ATS) for immunofluorescent analyses, labelled with GFP, cDcp-1 (grey) and DAPI (blue). The dashed lines outline GFP-positive cells. Scale bar, 50 μm.

**Fig. S3.**
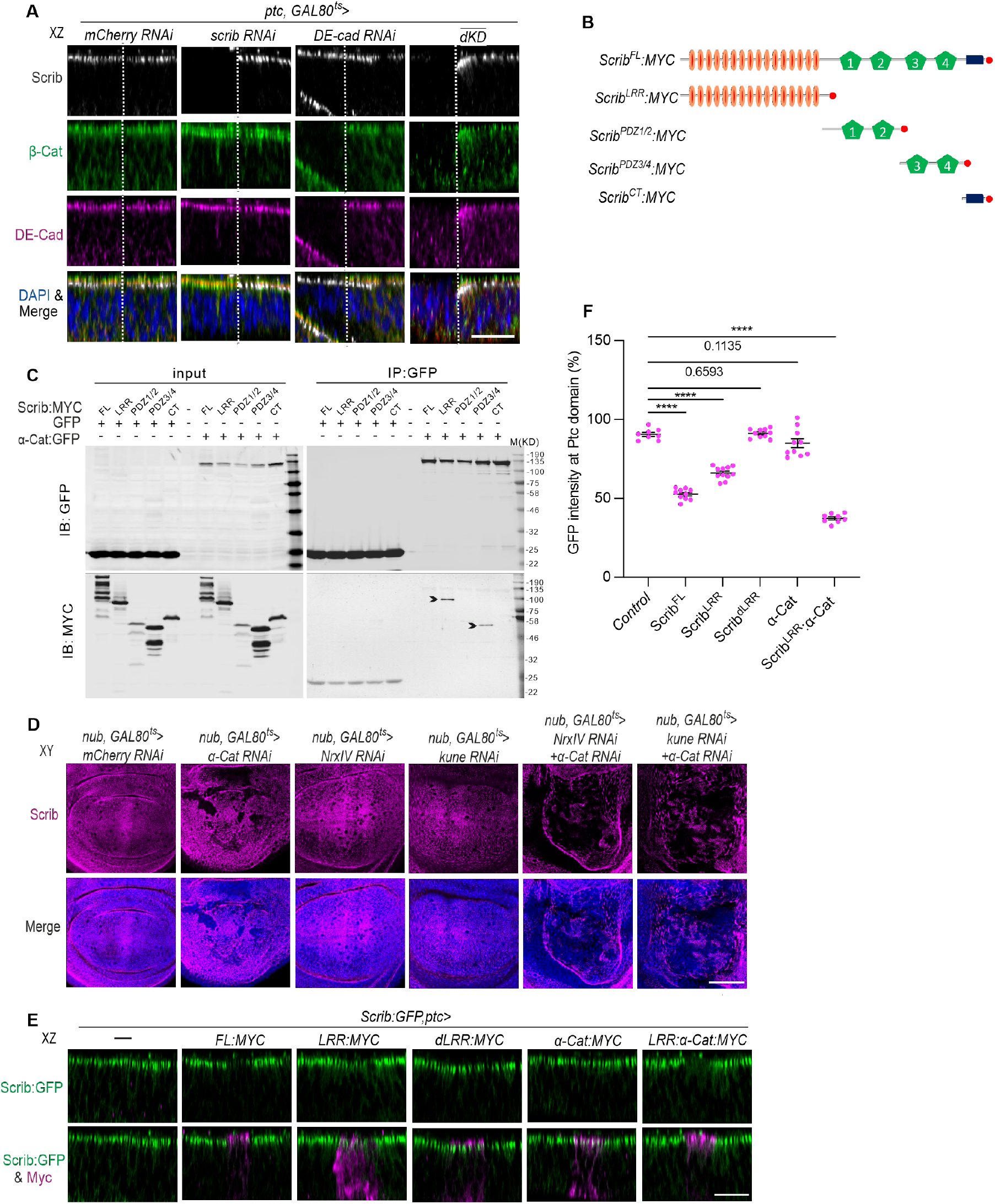
(A) Representative transverse section (XZ) images showing the wings with conditional expression of *mCherry* RNAi alone, *scrib* RNAi, *DE-cad RNAi* or *scrib* / *DE-cad RNAi* (dKD) driven by *ptc*-*GAL4*. Tissues were collected 2 days after temperature shift (ATS) for immunofluorescent analyses, labelled with Scrib (grey), β-Cat (green), DE-Cad (magenta) and DAPI (blue). The dashed lines outline the anterior-posterior boundary. Anterior is to the left. Anterior domain in the figure corresponds to *ptc* expression. (B) Schematics show the fragments of Scrib used in the immunoprecipitation assay (C) and immunostaining (E). (C) Western blotting showing Immunoprecipitation of different Scrib fragments tagged with MYC by α-Cat:GFP using GFP nanotrap. Arrowheads indicate LRR or PDZ3/4. (D) Representative coronal (XY) images showing the wings with conditional expression of *mCherry RNAi* alone, *α-Cat RNAi* and/ or *NrxⅣ RNAi, α-Cat RNAi* and/ or *kune RNAi* driven by *nub*-*GAL4*. Tissues were collected at 2 days after temperature shift (ATS) for immunofluorescent analyses, labelled with Scrib (magenta) and DAPI (blue). (E) Representative transverse section (XZ) images showing the wings with expression of *Scrib^FL^, Scrib^LRR^, Scrib^dLRR^*, α-Cat, or Scrib^LRR^:α-Cat driven by *ptc*-*GAL4* in Scrib:GFP animals. Tissues were collected for immunofluorescent analyses, labelled with Scrib:GFP (GFP) and Myc (magenta). (F) Quantification of relative Scrib:GFP intensity in (E) at the Ptc expressing domain. Sample sizes are 8 (control), 10 (Scrib^FL^), 11 (Scrib^LRR^), 10 (Scrib^dLRR^) 10 (α-Cat) and 8 (Scrib^LRR^:α-Cat). \*\*\*\**P* < 0.0001. Data are means ± 95 % confidence intervals (CIs). Statistical significance was calculated by the two-tailed *t*-test. Scale bars, 20 μm (A, E), 50 μm (D).

## Notes

### Competing Interest Statement

The authors have declared no competing interest.

